# Ubiquitination regulates granulostasis and DRiP accumulation in SGs under heat stress via the E3 ligase MKRN2

**DOI:** 10.1101/2025.10.15.682570

**Authors:** Emmanuel Amzallag, Yehuda M. Danino, Valentina Secco, Serena Carra, Eran Hornstein

## Abstract

Stress granules (SGs) are transient cytoplasmic biomolecular condensates that play a role in the cellular response to proteotoxic stress. It has been previously shown that ubiquitination regulates SG dynamics; however, the specific mechanisms by which ubiquitin affects SGs are not fully understood. Here, using proximity proteomics, we discover that the engagement of several E3 ubiquitin ligases in SGs is dependent on UBA1 activity. A detailed study of the E3 ubiquitin-protein ligase Makorin 2 (MKRN2) demonstrated that it is localized to SGs in a manner dependent on active ubiquitination. MKRN2 promotes both the proper formation of SGs and their disassembly following stress recovery, by preventing the accumulation of defective ribosomal products (DRiPs) within SGs. Therefore, MKRN2 is a novel regulator of SGs that mediates the maintenance of granulostasis, suggesting that the localization of a subset of E3 ligases into SGs is linked to their capacity to ubiquitinate target proteins.

## Introduction

Stress granules (SGs) are biomolecular condensates that form via a process of liquid-liquid phase separation (LLPS) of biomacromolecules (1). SG assembly is initiated by a variety of stressors, including heat stress, oxidative stress, osmotic stress, and ER stress, during which the cell activates the integrated stress response (ISR) (2). Phosphorylation of eIF2α initiates the ISR, resulting in polysome disassembly and global shutdown of translation. This releases mRNA molecules from ribosomes and engages a variety of RNA-binding proteins (RBPs) to form multivalent interactions with both RNA and with other proteins via RNA-binding domains (RBDs) and intrinsically disordered domains (IDD), respectively, resulting in the formation of discrete SGs. Upon recovery from stress, SGs rapidly disassemble, and their dissolution allows mRNAs to be redirected to translation (1).

However, SGs can fail to disassemble and persist for longer time, affecting the cells’ ability to restore translation, as well as the retro-translocation in the nucleus of RBPs (1). Several factors can delay SG disassembly including the accumulation of misfolded and damaged proteins and the impairment of the protein quality control system. The latter refers to molecular chaperones and protein degradation systems that cooperates to detect unfolded/misfolded proteins and promote their clearance (3). Amongst the misfolded and potentially toxic proteins that can accumulate inside SGs changing their dynamic liquid-like properties are disease-linked mutated RBPs such as TDP-43 or FUS or hnRNPA1 (4–6), as well as defective ribosomal products (DRiPs), which are truncated proteins released from disassembling polyribosomes. DRiPs are rapidly polyubiquitinated for degradation by the ubiquitin-proteasome system as well as by autophagy (7). Amongst the chaperones that participate in the maintenance of SG dynamics are the ATP-dependent chaperone complexes HSP70/HSPB8/BAG3 and VCP/UFD1L/PLAA. These chaperone complexes reduce DRiP accumulation inside SGs (8–10). Accordingly, the activity of HSP70, VCP, and also the autophagic machinery, thus helps surveil stress-induced DRiPs and maintain SGs functioning properly, in a process termed granulostasis (10).

Until now, the maintenance of granulostasis under stress has been largely associated with the activity of specific molecular chaperones, namely the HSP70/HSPB8/BAG3 and VCP/UFD1L/PLAA complexes (8,9). The ubiquitin-proteasome system (UPS) is a major component of cellular protein degradation and mediates the conjugation of ubiquitin onto target molecules marked for proteasomal degradation. While ubiquitination and SUMOylation, have been shown to impact SG dynamics as well (11–13) it is unknown if they regulate DRiPs accumulation in SGs.

Active ubiquitination is required for proper SG disassembly (11,14–17); however, the specific E3 ligases that maintain proper SG dynamics and granulostasis are only partially known. For example, the E3 ligase TRIM21 controls G3BP1 ubiquitination and SG condensation and the E3 ligase HECT regulates SG disassembly (16,17). However, whether specific E3 ligases regulate the clearance of DRiPs from SGs has not been yet explored.

Here, we report ubiquitination-dependent changes in SG proteomic composition that led to the identification of Makorin 2 (MKRN2) as an important SG-engaged E3 ligase that regulates SG dynamics and the sequestration of stress-induced DRiPs into SGs.

## Results

### Ubiquitination affects stress granule dynamics and composition

It was recently shown that active ubiquitination is required for proper SG disassembly (11,14,15,18). However, a detailed characterization of how SGs and the UPS interact has not been achieved. Using G3BP1-mCherry expressing U2OS cells (in which the endogenous G3BP1 and 2 genes were knocked out) (19) we studied the full SG assembly and disassembly cycle, induced by heat stress (HS; 43°C followed by stress recovery at 37°C; Fig. 1A,1B). Inhibition of the ubiquitin-activating enzyme UBA1 (ubiquitin-like modifier-activating enzyme 1) by TAK243 (1 μM) facilitated SG assembly, while delaying their dissolution (Fig. 1A-C, Supplementary fig. 1). Treatment with TAK243 coupled with HS caused SGs to nucleate and begin assembling earlier than in HS-treated cells (Fig.1B, 1C). SG disassembly following recovery from heat stress, at 37°C, was also delayed (Fig.1A, B, D, Supplementary fig. 1).

**Figure 1.**
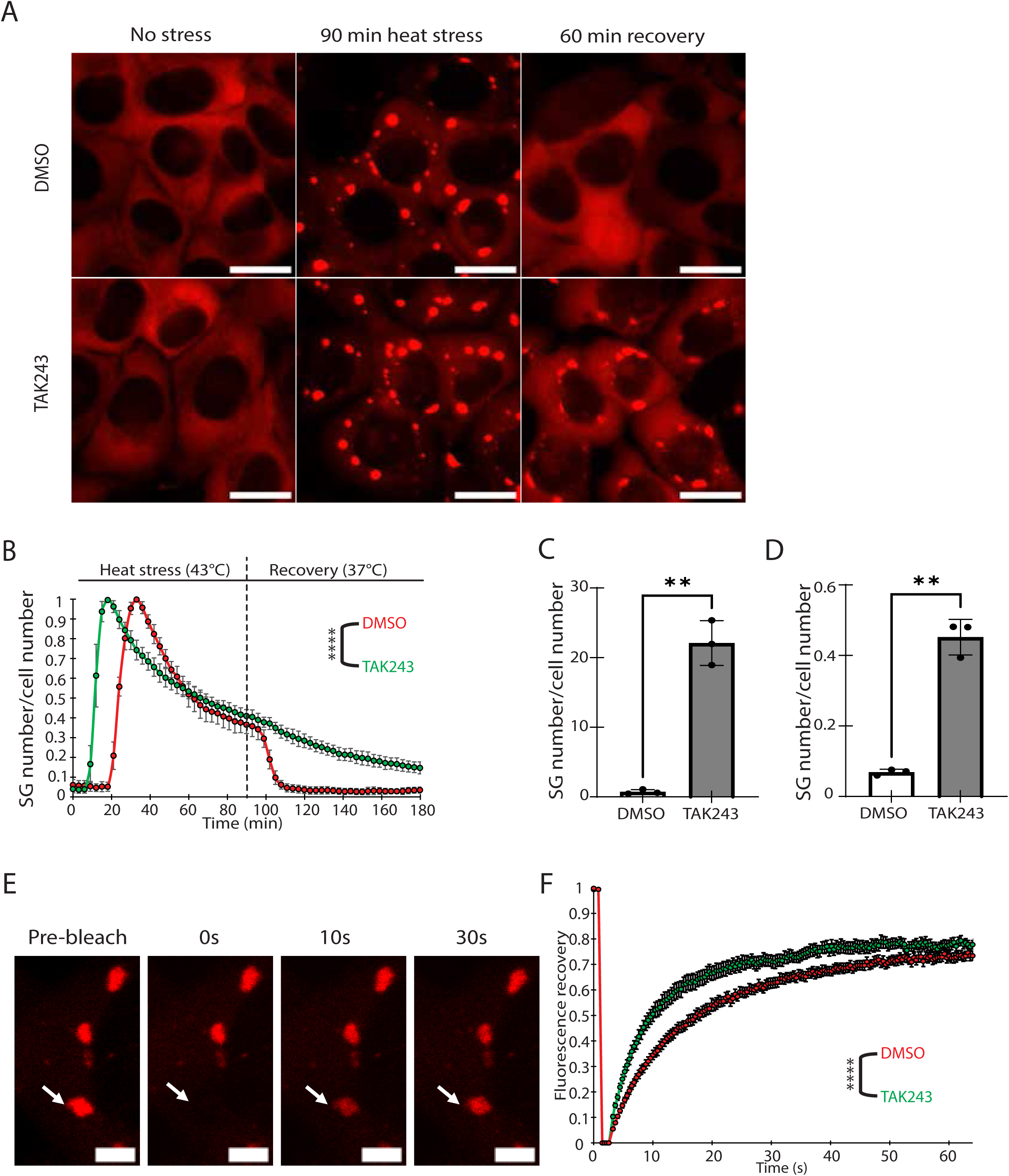
Ubiquitination affects stress granule dynamics and biophysical properties. **A.** Representative images of U2OS cells overexpressing G3BP1-mCherry with G3BP1/2 KO background, without (DMSO) or with TAK243 (1uM) after 90 min of heat stress at 43 °C (HS) or during recovery at 37 °C for 1h. Scale bar = 20uM. **B.** Quantification of SG number/cell number as a function of time. Data were normalized internally to the time point with maximal SG assembly. ∼1000 SG counted per experimental repeat. Nine fields per condition. Data from one experiment out of three experimental repeats (Supplementary fig. 1). Two-way ANOVA with repeated measures. **C.** Quantification of SG number/cell number at 15 min of HS. Average of three experimental repeats shown. Welch’s t-test. **D.** Quantification of SG number/cell number at 60 min recovery. Average of three experimental repeats shown. Welch’s t-test. **E.** A temporal image series from a FRAP experiment on G3BP1-mCherry U2OS cells under heat stress. Arrow indicates photobleached SG. Scale bar = 5uM. **F.** Quantification of fluorescence recovery after photobleaching in U2OS cells without or with TAK243. Data from one experimental repeat out of three is shown (Supplementary fig. 6). ∼20 SGs were counted per experiment. Two-way ANOVA with repeated measures. P value * <0.05; ** <0.01, ****<0.0001.

To determine whether active ubiquitination affects SG liquidity, we performed fluorescence recovery after photobleaching (FRAP) on cells under HS, following UBA1 inhibition. Inhibition of ubiquitination delayed fluorescence recovery dynamics of G3BP1-mCherry, relative to HS treated cells (Figure 1F, Supplementary fig. 6A). These observation indicate that ubiquitination is required for maintaining SG liquidity, consistent with previous reports (14).

To characterize the composition of SGs and identify players of the ubiquitination pathway that are involved in maintaining SG dynamics, we used APEX proximity labeling coupled to mass spectrometry (APEX-MS). G3BP1/2 KO U2OS cells expressing G3BP1-APEX were used to label SG proteins, while U2OS cells expressing nuclear export signal (NES)-APEX were used to label the surrounding cytoplasm). Heat stress was induced (43°C; 90 min), UBA1 was inhibited with TAK243, and proximity labeling was induced with biotin phenol (500 μM) and a short hydrogen peroxide pulse (1mM, 60 seconds). Biotin-labelled proteins were isolated with streptavidin magnetic beads and analyzed by mass spectrometry. Samples without biotin-phenol were used to identify and subtract non-APEX-derived biotinylation background. Peptide reads were mapped to proteins with the MaxQuant search engine (v1.6.6.0). Relative to the NES cytoplasmic control, 6200 proteins interacting with G3BP1-APEX were identified, including previously-reported SG proteins (Fig. 2A), providing confidence that our protocol effectively isolated SG proteomes. To understand which proteins are associated with SGs in a ubiquitination-dependent manner, we next compared the HS-induced SG proteomes without or with TAK243 (Fig. 2B, Supplementary fig. 2A,B). This provided a list of proteins either enriched (83 proteins) or depleted (215 proteins) from SGs upon inhibition of ubiquitination (Student’s T-test, FDR<0.05; Fig. 2C). Amongst the proteins that associated with SGs in a ubiquitin-dependent manner we report HSPA12, DNAJA1, and the VCP cofactors, UFD1L, NSFL1C, FAF1, and PLAA. Interestingly, these chaperones were previously implicated in promoting SG disassembly by preventing aberrant accumulation of stress-damaged proteins (9,14). Ubiquitin hydrolases and E3 ligases were also associated with SGs in a ubiquitination-dependent manner, including the RNA-binding ubiquitin ligases (RBULs) MKRN2, CNOT4, TRIM25, and ZNF598. We orthogonally validated the ubiquitin-dependent interaction of the most SG-depleted E3 ligase, MKRN2, by immunofluorescence in U2OS cells fixed after 90 min HS, without or with TAK243 (Fig. 2D, 2E). In agreement with APEX-MS results, endogenous MKRN2 exhibited decreased SG partitioning in response to TAK243 treatment (Fig. 2E).

**Figure 2.**
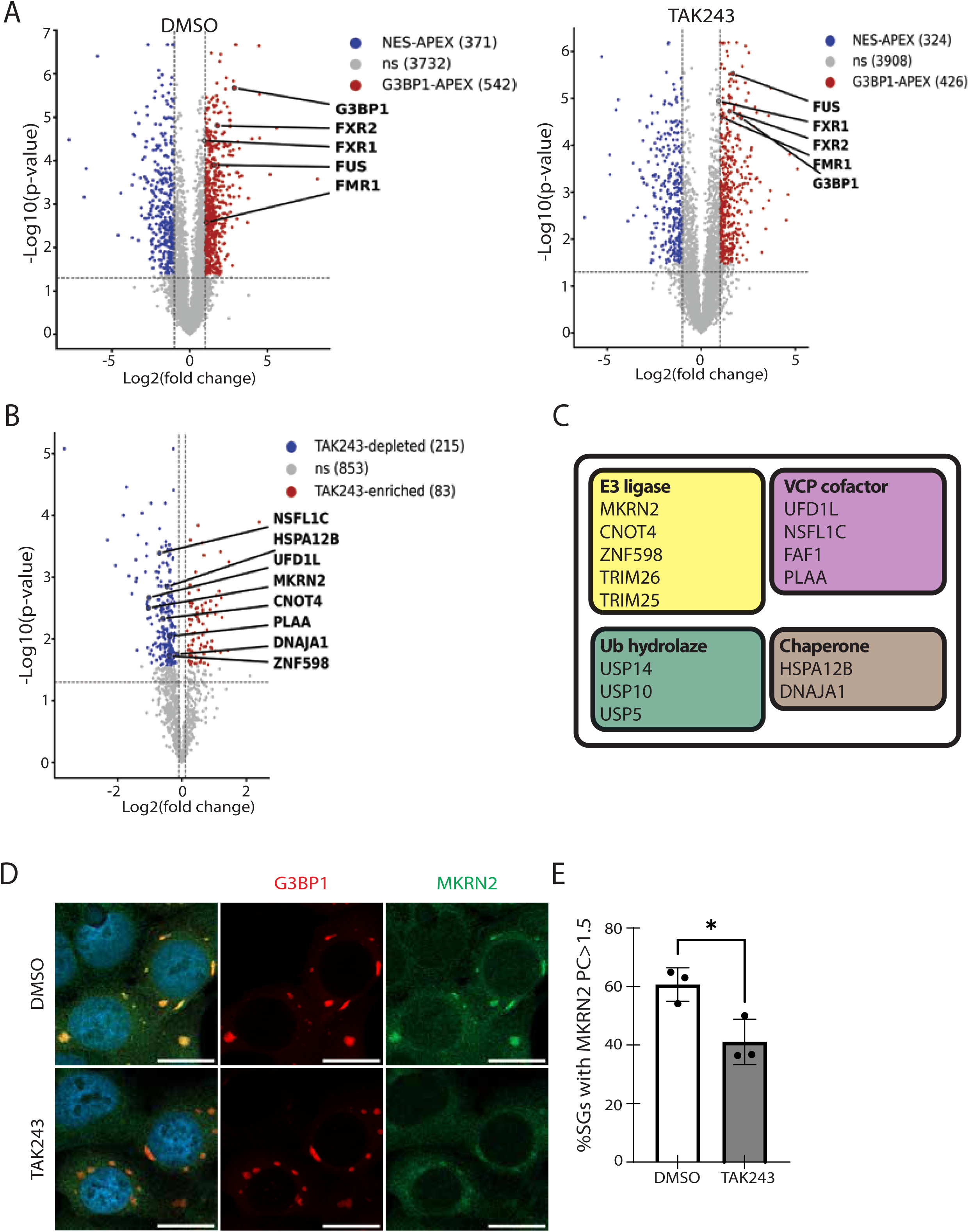
Ubiquitination affects stress granule proteomic composition. **A.** Volcano plot of proteins enriched at the proximity of G3BP1-APEX relative to NES-APEX cytoplasm control (x-axis = Log2 fold change) and statistical significance (y-axis = Log10 p-value). G3BP1/2 KO cells expressing G3BP1-APEX were used to label SG proteins. Induction media contained Biotin phenol, 500uM, and H2O2, 1mM. Labeling activity was quenched 60 seconds after H2O2 addition. **B.** Volcano plot displaying the ubiquitin-dependent enrichment in SG proteins. 83 proteins enriched and 215 proteins depleted from SGs following TAK243 treatment. (x-axis = LogFC; y-axis = log10 p-value). Several E3 ligases, ubiquitin hydrolases, and chaperones (shown in red) are depleted. **C.** Table showing proteostasis factors depleted from SGs upon inhibition of ubiquitination. **D.** Representative images of G3BP1-mCherry cells treated with TAK243 and stained for endogenous MKRN2. **E.** Quantification of the partition coefficient of MKRN2 in SGs in cells treated with DMSO or TAK243. The proportion of SGs with PC>1.5 is shown. Average of three experimental repeats, ∼1000 SGs per repeat. Welch’s two-sided t-test, p-value * = <0.05; **<0.01, ****<0.0001.

### The E3 ligase MKRN2 affects SG dynamics

While the human genome encodes only 2 E1s and ∼30 E2 enzymes, ∼600 E3 ligases are known, conferring substrate specificity in targeting specific proteins and cellular pathways (20). To determine whether ubiquitin-sensitive RBULs ligases control SG dynamics, we knocked down CNOT4, ZNF598, and MKRN2 in U2OS cells that were subjected to HS, followed by recovery (43 °C, 90 min -> 37 °C). Knockdown of CNOT4, ZNF598, or MKRN2 all impacted SG dynamics to varying degrees (Supplementary fig. 5A, B). MKRN2 depletion affected both SG assembly and disassembly, with cells forming smaller SGs that took longer to disassemble upon recovery (Figure 3,4). ZNF598 similarly affected SG size but had a smaller impact on disassembly. CNOT4 had a minimal effect on SG assembly, with a modest effect on their disassembly (Supplementary fig. 5A, B). Based on our mass spectrometry data, indicating MKRN2 as the most highly SG-depleted E3 upon TAK243 treatment and its strongest impact on SG dynamics, we focused mechanistic studies on MKRN2.

**Figure 3.**
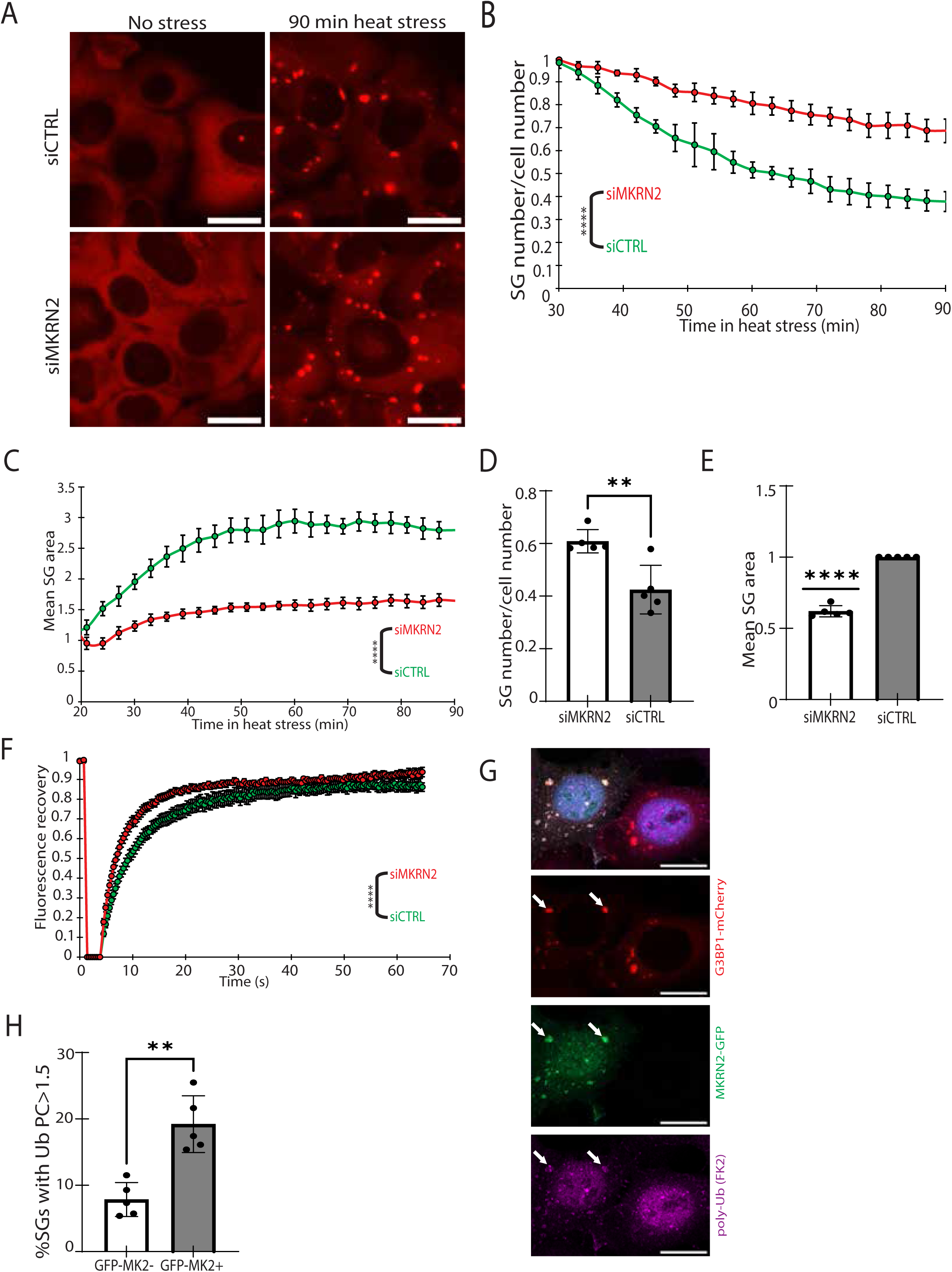
MKRN2 promotes SG assembly and increases SG ubiquitin content. **A.** Representative images of G3BP1-mCherry cells with siMKRN2 vs siCtrl before HS and at 90 min HS at 43 °C. **B.** Quantification of SG number/cell number from 30 min to 90 min HS at 43 °C. Data were normalized internally to the time point with maximal SG assembly. ∼1000 SG counted per experiment. Nine fields per condition. Data from one experimental repeat out of five independent experiments. Two-way ANOVA with repeated measures. **C.** Quantification of SG number/cell number at 90 min HS. Average of five experimental repeats shown. Welch’s t-test. **D.** Average SG area over the 90 min HS at 43 °C. Data from one experiment out of five experimental repeats. (Supplementary fig. 3). Two-way ANOVA with repeated measures. **E.** Average SG area at 90 min HS at 43 °C. Data were internally normalized to siCtrl. The average of five experimental repeats is shown. One-sample t-test. **F.** Quantification of fluorescence recovery after photobleaching in U2OS cells with either siMKRN2 or siCTRL. Data from one experimental repeat out of three is shown. (Supplementary fig. 6). ∼20 SGs were counted per experiment. Two-way ANOVA with repeated measures. **G.** Confocal micrographs of G3BP1-mCherry/MKRN2-GFP U2OS cells stained for poly-ubiquitin (FK2). **H.** FK2 partition coefficient quantification in SGs from either MKRN2-GFP+ vs MKRN2-GFP- cells within the same coverslip. The proportion of SGs with PC>1.5 is shown. Average of 5 experimental repeats shown. >1000 SGs per repeat. p-value * = <0.05; **<0.01, ****<0.0001.

**Figure 4.**
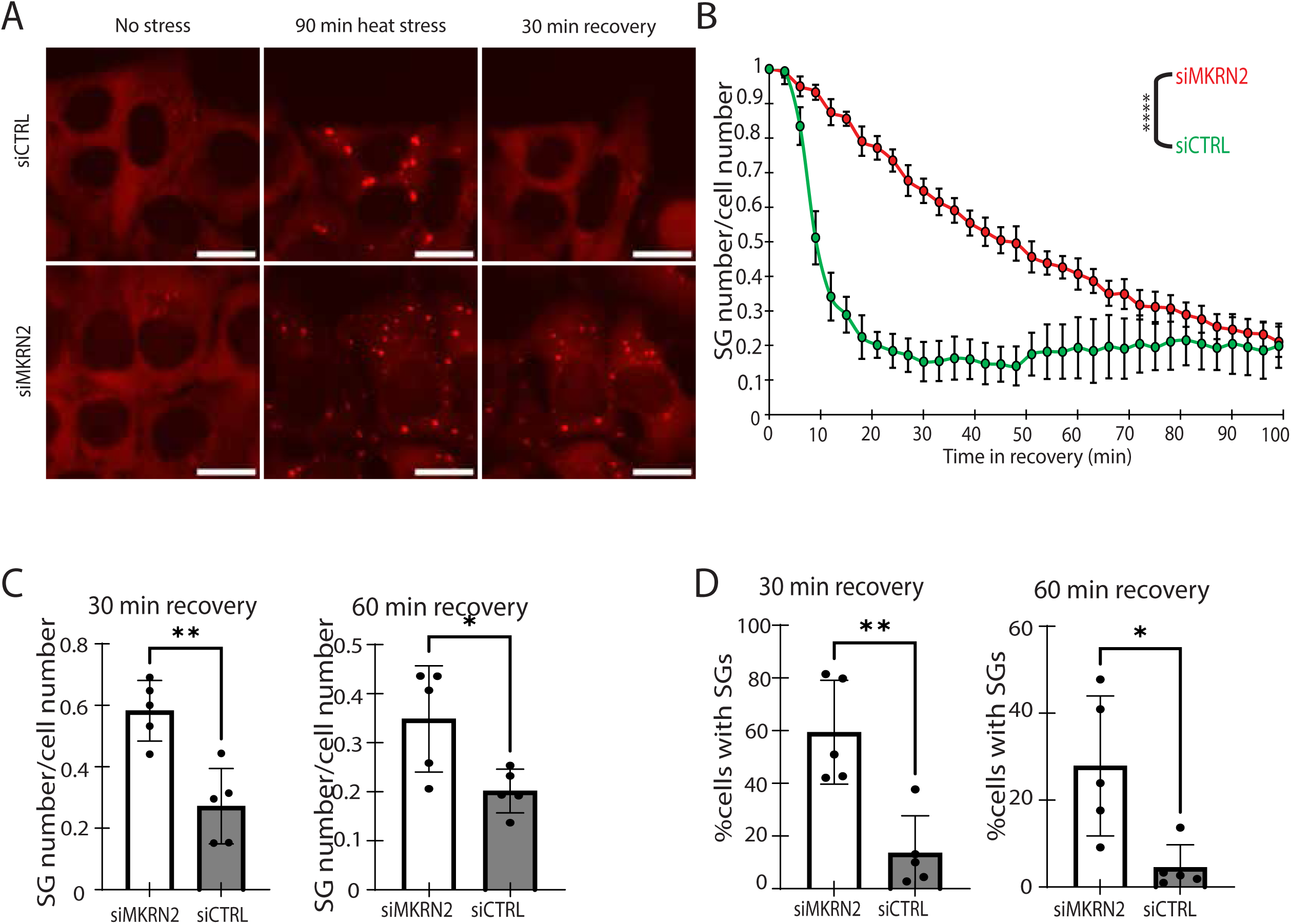
MKRN2 promotes SG disassembly. **A.** Representative images of G3BP1-mCherry cells with siMKRN2 vs siCtrl before HS, at 90 min HS at 43 °C and at 30 min recovery at 37 °C. **B.** Live imaging quantification of SG number/cell number during recovery at 37 °C. Data were internally normalized to 90 min of HS. 9 sites per condition. Data of one experiment out of five experimental repeats. Two-way ANOVA with repeated measures. **C.** Quantification of SG number/cell number at 30 min or 60 min recovery for siMKRN2 vs siCTRL. Average of five experimental repeats shown. Welch’s t test. **D.** Quantification of percent cells with SGs at 30 min or 60 min recovery for siMKRN2 vs siCTRL. Average of five experimental repeats shown. Welch’s t-test. p-value * = <0.05; **<0.01, ****<0.0001.

Using SG live imaging, we investigated the consequences of MKRN2 KD on SG dynamics under HS. SG assembly under heat stress typically involves a reduction in SG numbers over time, as smaller condensates coalesce to form larger but fewer SGs (8,21–23). The knockdown (KD) of MKRN2 resulted in a higher number of SGs at 90 min of HS (Fig. 3A, B, D) and the average size of SGs was almost half as small compared to the control (Fig. 3C, El Supplementary fig. 3). These data suggest that SGs fail to coalesce and mature upon MKRN2 KD. Having established that MKRN2 depletion affects SG number and size, we then measured their liquid-like properties by performing FRAP experiments, using the mobility of G3BP1 as readout. Consistently, FRAP analysis demonstrated that MKRN2 KD resulted in a slight increase in SG liquidity (Fig. 3F, Supplementary fig. 6B), that could be interpreted as associated to decreased coalescence into dense and mature SGs (23,24).

Then, to further test the impact of MKRN2 on SG dynamics, we overexpressed MKRN2–GFP and we observed that it was recruited to SG (Fig. 3G). We next quantified SG ubiquitin content in response to MKRN2-GFP overexpression, using an antibody that recognizes free, mono- and poly-ubiquitinated proteins (FK2). MKRN2-GFP increased the partitioning of ubiquitin into SGs, relative to neighboring cells that did not express MKRN2-GFP in the same polyclonal mosaic culture (Fig. 3G, 3H). Together these data indicate that MKRN2 regulates the assembly dynamics of SGs by promoting their coalescence during HS and can increase SG ubiquitin content.

We next tested the impact of MKRN2 on SG disassembly following stress removal. MKRN2 KD delayed the disassembly of SGs (Fig. 4A, 4B; Supplementary fig. 4). The effect was pronounced 30 and 60 minutes after cessation of HS (Fig. 4C, 4D). Therefore, MKRN2 is required for maintaining proper SG dynamics under HS.

### MKRN2 regulates accumulation of DRiPs in SGs

DRiPs are truncated peptides that are prone to aggregation and DRiPs ubiquitination affects their solubility and clearance (7,8,25). Proteostasis impairment causes aberrant DRiP accumulation inside SGs and impairs SG disassembly. This suggests the idea that MKRN2 may ubiquitinate a subset of DRiPs, avoiding their accumulation inside SGs and facilitating their disposal. To test the idea that ubiquitin impacts a subset of DRiPs in SGs, we induced SG formation by HS in absence or presence of the UBA1 inhibitor TAK243. Concomitantly we treated the cells with O-propargyl puromycin (OP-puro, 25uM), which is incorporated into nascent polypeptide chains, causing polysome disassembly and the emergence of DRiPs. We observed increased partitioning of OP-puro labelled DRiPs in SGs upon inhibition of ubiquitination relative to control (Fig. 5A, 5B). These data suggest that active ubiquitination prevents the accumulation of DRiPs inside SGs. To understand whether a fraction of DRiPs is ubiquitinated by MKRN2, we then performed a similar analysis in proficient and MKRN2-deficient cells. We measured increased partitioning of OP-puro labelled DRiPs in SGs upon knockdown of MKRN2, relative to control (Fig. 5C, 5D). Therefore, we conclude that MKRN2 ubiquitinates a subset of DRiPs, avoiding their accumulation inside SGs.

**Figure 5.**
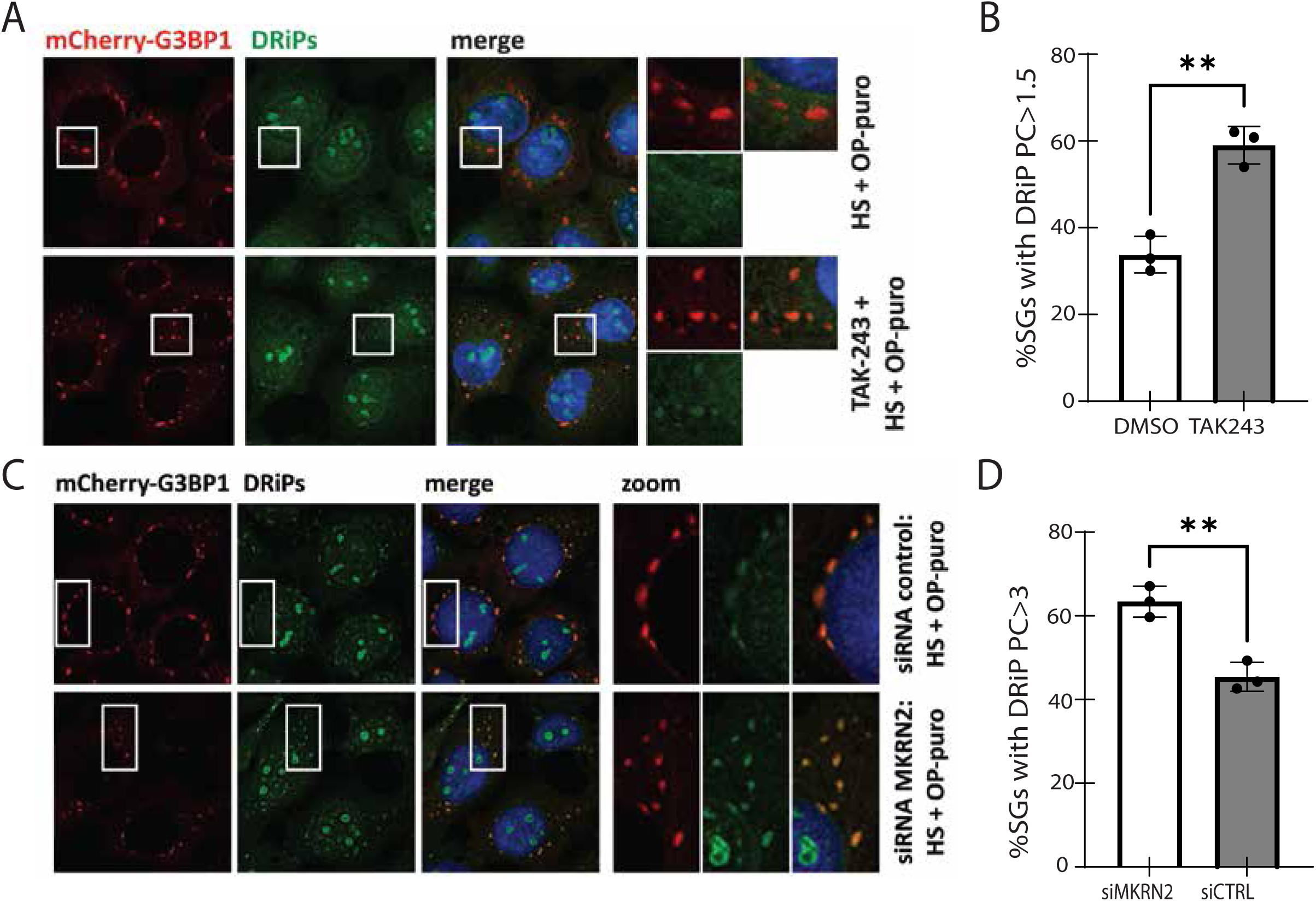
Ubiquitination is required for preventing DRiP accumulation in SGs. **A.** Representative images of DMSO vs TAK243 G3BP1-mCherry U2OS cells treated with 25 μM OP-puro to label DRiPs and heat-stressed at 43 °C for 60 min to induce SGs. **B.** Quantification of the partition coefficient of DRiPs in SGs from cells treated with either DMSO or TAK243. The proportion of SGs with PC>1.5 is shown. Welch’s t-test. **C.** Representative images of siMKRN2 vs siCTRL G3BP1-mCherry U2OS cells treated with 25 μM OP-puro to label DRiPs and heat-stressed at 43 °C. **D.** Quantification of the partition coefficient of DRiPs in SGs from cells treated with either siMKRN2 or siCTRL. The proportion of SGs with PC>1.5 is shown. Welch’s t-test. **<0.01

Together our study provides evidence for a new role for ubiquitination and for the E3 ligase MKRN2 in the control of SG dynamics and DRiPs turnover.

## Discussion

In this study, we characterize the impact of ubiquitination on SG proteomic composition during heat stress. Using APEX proximity labeling and mass spectrometry, we demonstrate that ubiquitination regulates SG functions under heat stress by controlling the recruitment of several molecular regulators of proteostasis into SGs.

Specifically, ubiquitination regulates the localization of several proteostasis-related proteins to SGs, including four VCP cofactors (UFD1L, PLAA, NSFL1C, and FAF1), two of which (UFD1L and PLAA) have been shown to regulate SG dynamics and sequestration of DRiPs (8). This suggests that ubiquitin-binding VCP cofactors are recruited to SGs through their interaction with ubiquitinated proteins within these condensates. Furthermore, the localization of the heat shock protein HSP70 (HSPA12B) and its co-chaperone DNAJA1 to SGs is also regulated by ubiquitination. This finding is consistent with similar behavior observed under sodium arsenite stress (17) and is likely to influence the accumulation of DRiPs in SGs (9).

Several E3 ubiquitin ligases were depleted from SGs in the absence of UBA1 activity, namely MKRN2, ZNF598, CNOT4, TRIM25, and TRIM26, whereas the E3 RNF20 was enriched in SGs. A detailed study of the E3 ligase MKRN2 demonstrated ubiquitination-dependent partitioning of endogenous MKRN2 into SGs. This suggests that the localization of a subset of E3 ligases into SGs is linked to their capacity to ubiquitinate target proteins.

Pan et al. reported enrichment of several HECT-domain E3 ligases in SGs following inhibition of ubiquitination by TAK243 (17), suggesting that this process is specific and potentially regulated in a stress- or protein-specific manner. SG E3 ligases that depend on UBA1 activity are RBULs, and their capacity to bind RNA may be particularly intriguing for SG E3s.

Endogenous and ectopically expressed MKRN2 were both localized to SGs. MKRN2 has a functional role in regulating SG dynamics under heat stress, and its knockdown results in more numerous and smaller SGs, as well as a significant delay in SG disassembly following recovery from stress.

We also show that UBA1 activity and MKRN2 regulates a subset of DRiP homeostasis in SGs. Thus, ubiquitination is required for preventing excessive DRiPs accumulation in SGs under heat stress. We further show that the RBUL MKRN2 similarly prevents accumulation of DRiPs in SGs suggesting that it may execute ubiquitin-mediated extraction of DRiPs from SGs. We note that our experiments cannot delineate whether ubiquitination of DRiPs by MKRN2 occurs inside SGs or in the cytoplasm. However, the finding that MKRN2 is enriched in SGs under stress coupled with the fact that its overexpression increases SG ubiquitin content, suggests that MKRN2 controls DRiP clearance from SGs via polyubiquitination of SG-localized substrates. In summary, MKRN2 is the first E3 ligase shown to play a role in DRiP extraction from SGs. The phenotype of smaller, more numerous SGs concomitant with increased DRiP partitioning, also reported upon inhibition of autophagy and VCP function (8), suggests that MKRN2 may act in the same pathway.

We note that the SG disassembly phenotype upon MKRN2 knockdown is less pronounced than that observed with TAK243 treatment. Therefore, MKRN2 may target a subset of DRiPs. consistent with the role of additional SG-localized E3 ligases, such as ZNF598 or CNOT4, that regulate SG dynamics. Overall, our results support the conclusion that proper SG dynamics depend on clearance of DRiPs from these condensates and portray MKRN2 as a novel proteostasis factor regulating this process.

Limitations: The direct MKRN2 substrates and ubiquitin-chain types (K63/K48) are currently unknown. In addition, it is interesting to understand if DRiP solubility is dependent on ubiquitination or on specific E3 ligases, such as MKRN2.

In sum, global ubiquitination promotes the recruitment of key SG-engaged chaperones, including HSP70 and VCP, as well as a subset of E3 ubiquitin ligases. We identify the E3 ligase MKRN2 as a novel regulator of SG dynamics and proteostatic function. Clearance of DRiPs from SGs requires active ubiquitination and is mediated in part by MKRN2. These insights refine our understanding of how SGs coordinate the triage of stress-damaged proteins with condensate dynamics during proteotoxic stress. Ultimately, defining ubiquitination-dependent regulation of SG composition and function may illuminate new therapeutic avenues for restoring proteostasis in diseases that involve protein aggregation.

## Methods

### Mammalian cell culture

U2OS cells were cultured in growth media consisting of high glucose DMEM (Sartorius 01-052- 1A), 10% FBS (Sigma F7524) and 1% penicillin-streptomycin-amphotericin B (Sartorius 03-033- 1B) in a 37°C incubator supplemented with 5% CO2. To induce heat stress, cells were placed into a dedicated incubator set to 43°C supplemented with 5% CO2. For UBA1 inhibition, cells were pre-treated with 1uM TAK243 (Med Chem Express; Cat# HY-100487) prior to heat stress. For MKRN2 KD, cells were transfected with MKRN2 siRNA (Dharmacon siGENOME SMARTpool; M-006960-01-0005) using Lipofectamine RNAiMAX (Cat#: 13778075) and cultured for 72 hours before analysis.

### Molecular cloning

Molecular cloning of MKRN2-GFP was performed using In-Fusion® Snap Assembly (Cat# 638947; Takara Clontech) and inserted into pcDNA4.0 vector with puromycin resistance. G3BP1-mCherry cells were transfected with Lipofectamine 2000 (Thermo Fisher Scientific, Cat# 11668027) cultured with 5ng/ml puromycin for 7 days.

### Microscopy

SG live imaging was performed in a Zeiss CD7 widefield microscope equipped with an Axiocam 702 CMOS camera using a 20X air objective with an additional 2X post-magnification (effective magnification of 40X, excitation wavelength 590nm). G3BP1-mCherry cells were seeded into optical 24-well plates (Cellvis P24-1.5H-N) and placed into the heated microscope incubator chamber (37°C, 5% CO2). After one cycle of imaging, the microscope incubator temperature was increased to 43°C. After 90 min of heat stress, the temperature was lowered back down to 37°C to image SG disassembly. SG number and area was analysed using the surfaces feature in Imaris software. Immunofluorescent staining was performed on 12mm round glass coverslips inside 24-well plates. Immunofluorescent images were captured in a Zeiss LSM900 confocal microscope using a 63X oil objective lens.

### FRAP

G3BP1-mCherry cells were seeded into 3.5mm dishes with a central coverslip (MatTek P35GC- 1.5-14-C). Cells were imaged in a Leica SP8 STED microscope using a 100X objective. To induce heat stress, the microscope incubator temperature was increased to 43°C and an objective heater was attached to the objective and heated to 43°C. Cells were left in the heated microscope incubator chamber for 60 minutes, and FRAP was performed for 30 minutes thereafter. To photobleach SGs, a 561nm laser set to max power was used to bleach entire SGs. In order to monitor fluorescence recovery, images were collected every 400ms for 63 seconds. Fluorescence recovery data was collected using the Leica LasX software. Analysis corrected for background fluorescence intensity and normalized to the pre-bleach SG intensity to calculate the fraction of fluorescence recovery at each time point.

### APEX proximity labelling

G3BP1-APEX or NES-APEX U2OS (Marmor-Kollet *et al.*) cells were seeded into T125 flasks with 50ng/ml tetracycline (Sigma T7660) and allowed to incubate for 24h before performing APEX experiment. Four technical repeats were used for each condition. Following pre-treatment with TAK243, cells were supplemented with 500uM Biotin-Tyramide (Iris Biotech 41994-02-9) for 90 minutes to allow for labelling activity. Labelling was then induced by addition of H2O2 (JT Baker 2186-01) for 60 seconds. APEX activity was extinguished with quenching solution (10mM sodium azide (Sigma cat number S2002), 10mM sodium ascorbate (Alfa Aesar A17759), 5mM Trolox (Sigma 238813) in PBS. Cells were then scraped in PBS, centrifuged at 3000xg for 10 minutes at 4°C and the cell pellet was lysed in ice-cold RIPA buffer (50mM Tris-HCl pH8, 150mM NaCl, 1% NP-40, 0.5% sodium deoxycholate, 0.1% SDS) supplemented with cOmplete Protease Inhibitor Cocktail (Roche, 4693116001) and PhosSTOP (Roche, 4906837001). Lysates were centrifuged at 15000xg for 10 minutes at 4°C and protein concentration was quantified using Bradford assay (Bio-Rad 5000006). Streptavidin magnetic beads (Thermo Fisher cat number 88817) were incubated with the protein extracts at a ratio of 100ul beads per 500ug protein extract overnight at 4°C with rotation.

### Liquid chromatography and mass spectrometry

Samples were subjected to on-bead tryptic digestion. The resulting peptides were analysed using nanoflow liquid chromatography (nanoAcquity) coupled to high resolution, high mass accuracy mass spectrometry (Q-Exactive Plus). Each sample was analysed on the instrument separately in a random order in discovery mode. Raw data were processed with MaxQuant v1.6.6.0. The data were searched with the Andromeda search engine against the human proteome database, appended with common lab protein contaminants and the default modifications. Quantification was based on the LFQ method, based on unique/all peptides.

### Proteomics statistical analysis

ProteinGroups output table from MaxQuant was imported into Perseus for analysis. First, contaminants were removed. Raw intensity values were log2 transformed and missing values imputed for proteins with at least four valid values using an artificial normal distribution with a downshift of 1.8 standard deviations and a width of 0.3 of the original ratio distribution. To identify proteins non-specifically bound to streptavidin beads, student’s t test with FDR <=0.05 was used to compare each sample to respective no BP control samples. These non-specific binders were filtered out and proteins were then compared between G3BP1-APEX and NES-APEX baits. LFQ values were log2 transformed and missing values were imputed for proteins with at least two valid values in at least one group using an artificial normal distribution with a downshift of 1.8 standard deviations and a width of 0.3 of the original ratio distribution. Proteins were then compared between the G3BP1-APEX bait to the NES-APEX bait using student’s t test with FDR <=0.05 to determine which proteins were significantly enriched in proximity to G3BP1 compared to NES. G3BP1-APEX values were then normalized by the mean of their corresponding NES-APEX values. The NES-normalized values were then compared between the TAK243 vs DMSO treated samples using student’s t test with FDR <=0.05.

### Statistical analysis

Statistical analysis of live imaging data was done using repeated-measures ANOVA test with contrast analysis in R. Pairwise comparisons were done using Welch’s t test in Graphpad Prism. Statistical p values < 0.05 were considered significant.

### Labeling of nascent peptides with OP-puro and high content imaging-based assay

Cells were grown on polylysine-coated glass coverslip. Newly synthesized proteins were labelled by incubating the cells with 25 mM O-Propargyl-puromycin (OP-puro) for the indicated time points and CuAAC detection of OP-puro labelled DRiPs was performed as previously described, using Alexa Fluor 594 azide (A10270, Life Technologies) (PMID: 22160674). Cells were next processed for immunofluorescence microscopy. Blocking and incubation with primary and secondary antibodies were performed in PBS containing 3% BSA and 0.1% Triton X-100. DRiPs enrichment inside SGs was measured using the Scan^R Analysis software (Olympus).

SGs were segmented based on G3BP1 signal using edge detection algorithm. The mean fluorescence intensity of OP-puro labelled DRiPs was detected in each detected SG and was additionally measured in an area surrounding each SG. The relative enrichment of OP-puro labelled DRiPs in individual SGs was calculated as a ratio of mean fluorescence intensity inside the SG divided by mean intensity in the region surrounding the SG, as previously described (PMID: 27570075).

## Contributions statement

E.H. and E.A. oversaw the conceptualization of this study. E.A. and Y.M.D. performed the APEX proteomics experiment. Mass spectrometry was performed at the de Botton Institute for Protein Profiling from the Nancy and Stephen Grand Israel National Center for Personalized Medicine (G-INCPM). E.H. and E.A. designed the study and had unrestricted access to all data. E.A. performed and analyzed all live imaging experiments. E.A. performed immunofluorescence staining, microscopy and analysis of ubiquitin and MKRN2. E.A. performed all FRAP experiments. V.S. and S.C. performed the O-propargyl puromycin fluorescent staining, microscopy and analysis of DRiP partitioning in stress granules. E.A. and E.H participated in drafting the initial manuscript draft and E.H., E.A. and S.C. reviewed and revised the draft. All authors agreed to submit the manuscript, read and approved the final draft, and take full responsibility for its content, including the accuracy of the data and its statistical analysis.

## Acknowledgments

Eran Hornstein is the Mondry Family Professorial Chair and Head of the Nella and Leon Benoziyo Centre for Neurological Diseases and of the Andi and Larry Wolfe Centre for Neuroimmunology and Neuromodulation. The authors thank Dr. Yishai Levin, Dr. Alon Savidor, Corine Katina at the de Botton Institute for Protein Profiling from the Nancy and Stephen Grand Israel National Center for Personalized Medicine (G-INCPM) for performing mass spectrometry. The authors thank Yoseph Addadi, Inna Goliand and Tatiana Smirnova from the de Picciotto Cancer Cell Observatory In memory of Wolfgang and Ruth Lesser of the Moross Integrated Cancer Center in the department of Life Science Core Facilities, Weizmann Institute of Science for help with performing live imaging and FRAP. The authors thank Prof. Itay Koren for his advice on our study. The authors thank CIGS (University of Modena microscopy facility) and Dr. Jonathan Vinet for technical support for confocal microscopy image acquisition and analysis. Special thanks to Iddo Magen, Yahel Cohen and Aviad Siany from the Hornstein Lab for fruitful discussion and advice.

---

**Supplementary figure 1.**
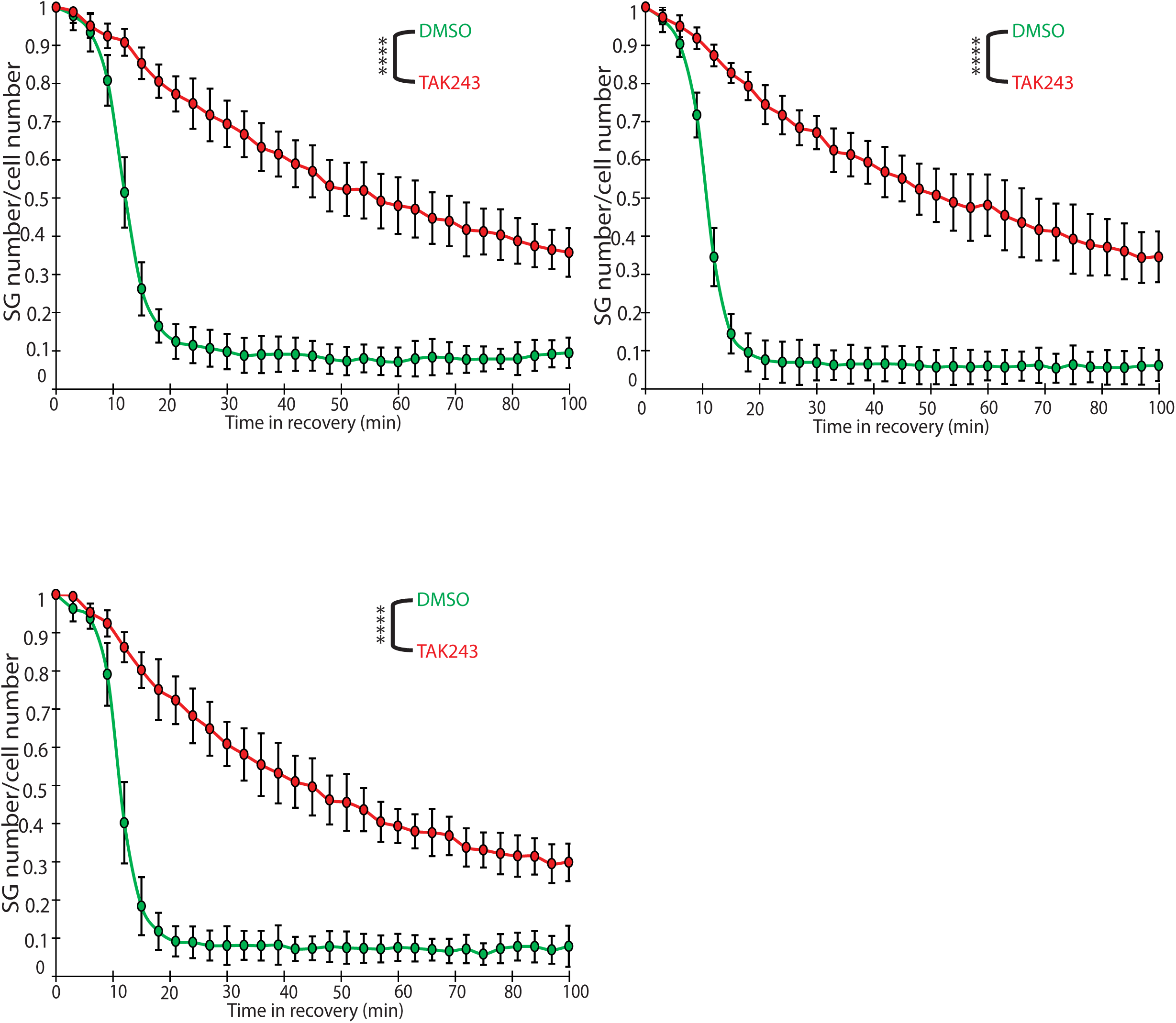
SG disassembly live imaging repeats with TAK243. Live imaging quantification of SG number/cell number during recovery at 37°C. Data were internally normalized to 90 min of HS. 9 sites per condition. Three experimental repeats shown. Two-way ANOVA with repeated measures. ****<0.0001.

**Supplementary figure 2.**
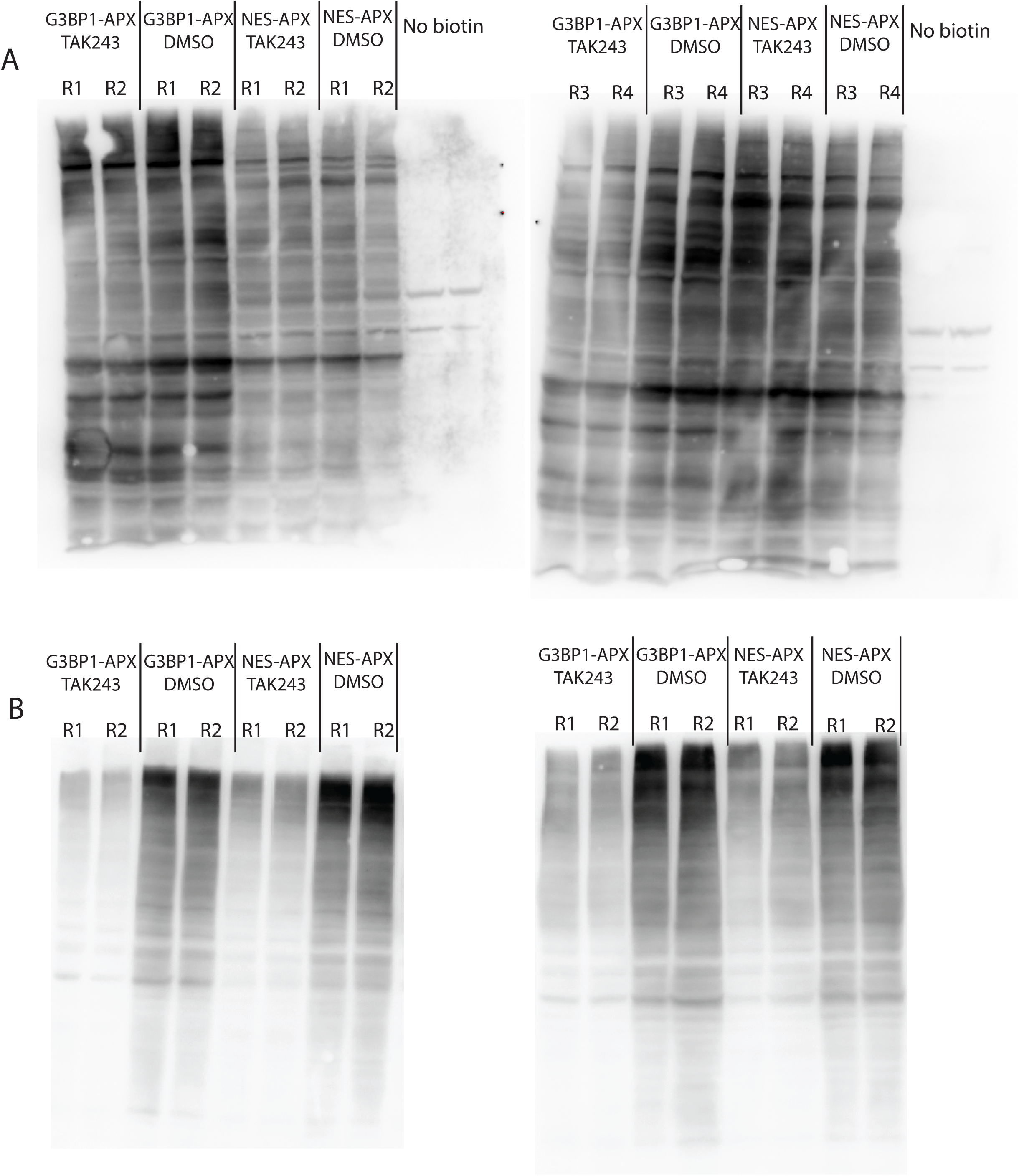
APEX samples. **A.** Anti-Biotin Western blot for samples used in APEX proteomics experiment. **B.** Ant-ubiquitin (FK2) Western blot for samples used in APEX proteomics experiment.

**Supplementary figure 3:**
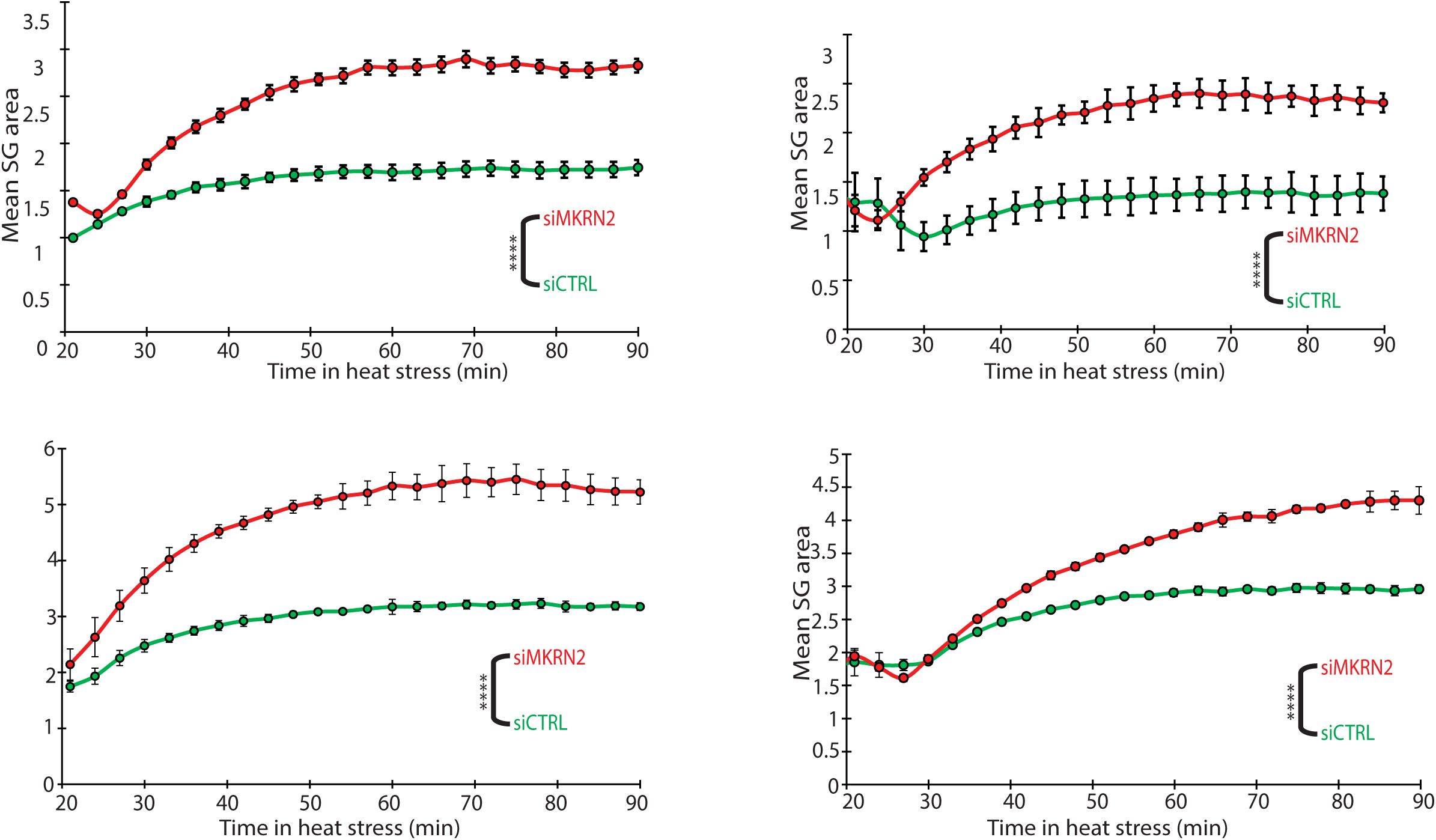
SG assembly live imaging repeats with MKRN2 KD. Live imaging quantification of SG area during HS at 43°C. 9-12 sites per condition. Four experimental repeats shown (fifth repeat shown in Fig. 3C). Two-way ANOVA with repeated measures. ****<0.0001.

**Supplementary figure 4.**
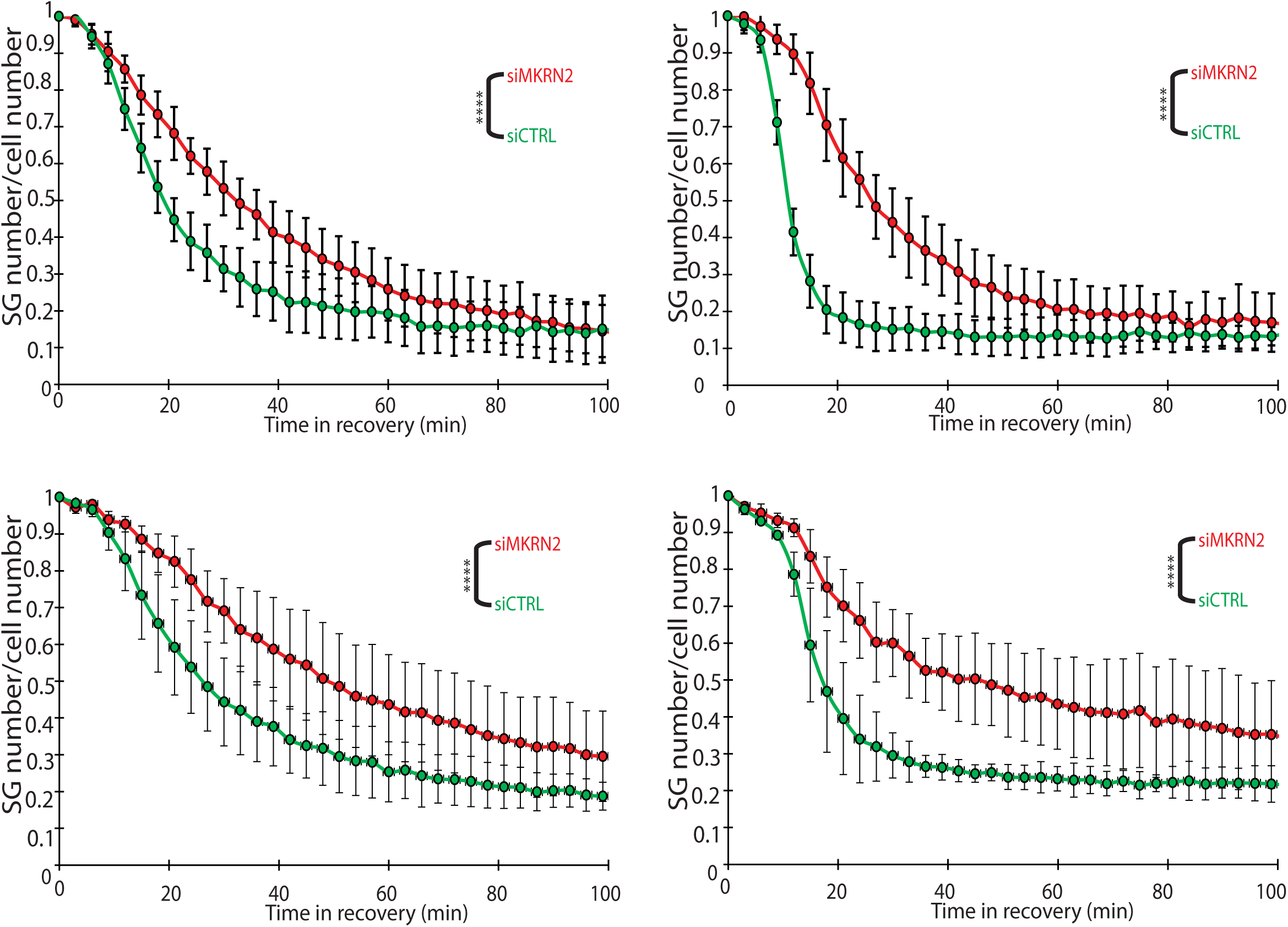
SG disassembly live imaging repeats with MKRN2 KD. Live imaging quantification of SG number/cell number during recovery at 37°C. Data were internally normalized to 90 min of HS. 9-12 sites per condition. Four experimental repeats shown (fifth repeat shown in Fig. 4B). Two-way ANOVA with repeated measures. ****<0.0001.

**Supplementary figure 5.**
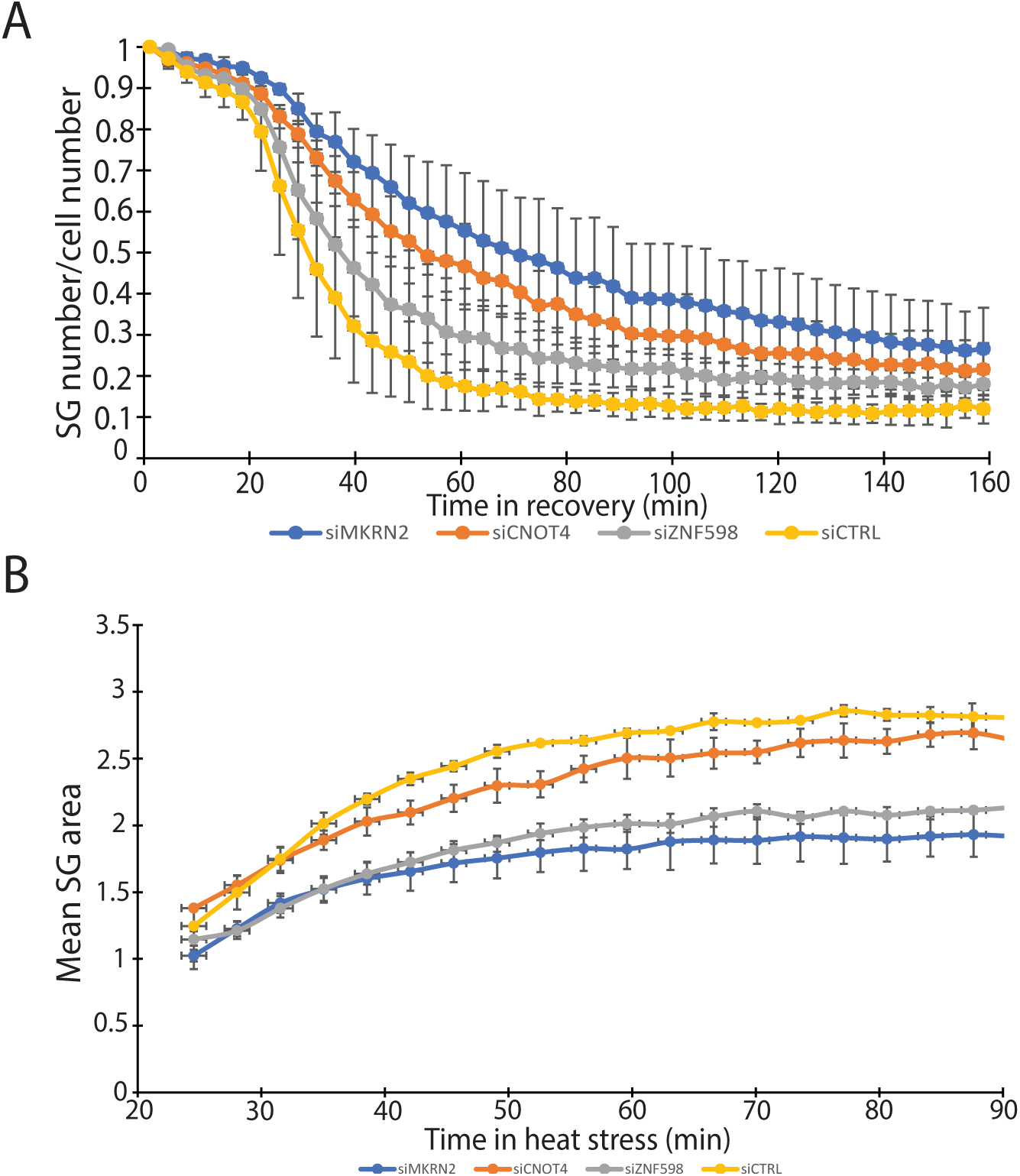
Live imaging with MKRN2, ZNF598 and CNOT4. **A.** SG number/cell number during stress recovery (normalized to 90 min HS). **B.** Average SG area over time in HS.

**Supplementary figure 6:**
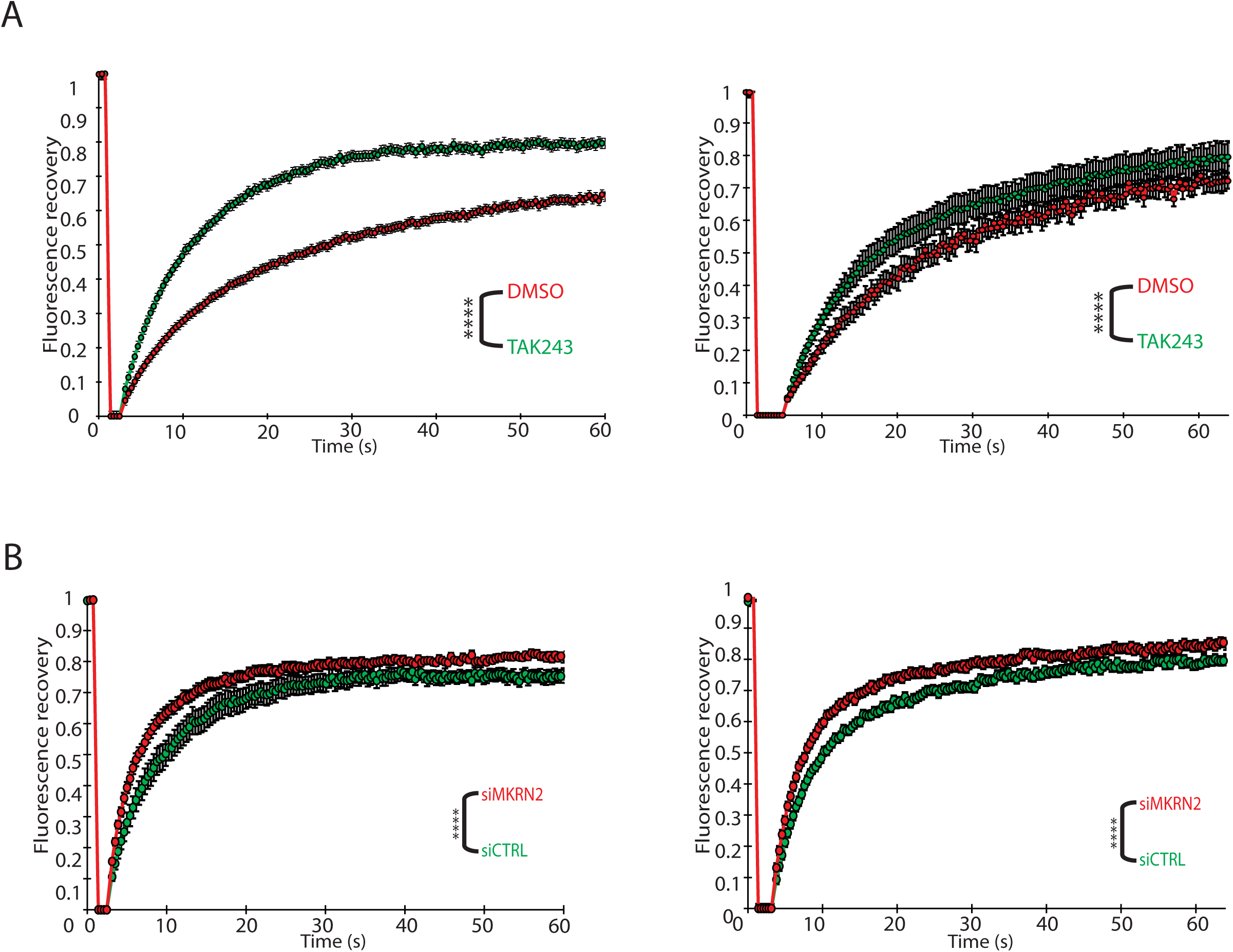
FRAP repeats. **A.** Quantification of fluorescence recovery after photobleaching in TAK243 vs control cells. Two experimental repats shown (third repeat in Fig. 1F). ∼20 SGs per repeat. **B.** Quantification of fluorescence recovery after photobleaching in TAK243 vs control cells. Two experimental repats shown (third repeat in Fig. 3F). ∼20 SGs per repeat. Two way ANOVA with repeated measures. ****<0.0001.

## References

1. Protter DSW, Parker R. Principles and properties of stress granules. Trends Cell Biol. 2016 Sept;26(9):668–79.

2. Bond S, Lopez-Lloreda C, Gannon PJ, Akay-Espinoza C, Jordan-Sciutto KL. The integrated stress response and phosphorylated eukaryotic initiation factor 2α in neurodegeneration. J Neuropathol Exp Neurol. 2020 Feb 1;79(2):123–43.

3. Hipp MS, Kasturi P, Hartl FU. The proteostasis network and its decline in ageing. Nat Rev Mol Cell Biol. 2019 July;20(7):421–35.

4. Gordon D, Dafinca R, Scaber J, Alegre-Abarrategui J, Farrimond L, Scott C, et al. Single-copy expression of an amyotrophic lateral sclerosis-linked TDP-43 mutation (M337V) in BAC transgenic mice leads to altered stress granule dynamics and progressive motor dysfunction. Neurobiol Dis. 2019 Jan;121:148–62.

5. Baron DM, Kaushansky LJ, Ward CL, Sama RRK, Chian R-J, Boggio KJ, et al. Amyotrophic lateral sclerosis-linked FUS/TLS alters stress granule assembly and dynamics. Mol Neurodegener. 2013 Aug 31;8(1):30.

6. Gui X, Luo F, Li Y, Zhou H, Qin Z, Liu Z, et al. Structural basis for reversible amyloids of hnRNPA1 elucidates their role in stress granule assembly. Nat Commun. 2019 May 1;10(1):2006.

7. Schubert U, Antón LC, Gibbs J, Norbury CC, Yewdell JW, Bennink JR. Rapid degradation of a large fraction of newly synthesized proteins by proteasomes. Nature. 2000 Apr 13;404(6779):770–4.

8. Seguin SJ, Morelli FF, Vinet J, Amore D, De Biasi S, Poletti A, et al. Inhibition of autophagy, lysosome and VCP function impairs stress granule assembly. Cell Death Differ. 2014 Dec;21(12):1838–51.

9. Ganassi M, Mateju D, Bigi I, Mediani L, Poser I, Lee HO, et al. A Surveillance Function of the HSPB8-BAG3-HSP70 Chaperone Complex Ensures Stress Granule Integrity and Dynamism. Mol Cell. 2016 Sept 1;63(5):796–810.

10. Alberti S, Mateju D, Mediani L, Carra S. Granulostasis: Protein quality control of RNP granules. Front Mol Neurosci. 2017 Mar 27;10:84.

11. Maxwell BA, Gwon Y, Mishra A, Peng J, Nakamura H, Zhang K, et al. Ubiquitination is essential for recovery of cellular activities after heat shock. Science. 2021 June 25;372(6549):eabc3593.

12. Marmor-Kollet H, Siany A, Kedersha N, Knafo N, Rivkin N, Danino YM, et al. Spatiotemporal Proteomic Analysis of Stress Granule Disassembly Using APEX Reveals Regulation by SUMOylation and Links to ALS Pathogenesis. Mol Cell. 2020 Dec 3;80(5):876–891.e6.

13. Verde EM, Antoniani F, Mediani L, Secco V, Crotti S, Ferrara MC, et al. SUMO2/3 conjugation of TDP-43 protects against aggregation. Sci Adv. 2025 Feb 21;11(8):eadq2475.

14. Gwon Y, Maxwell BA, Kolaitis R-M, Zhang P, Kim HJ, Taylor JP. Ubiquitination of G3BP1 mediates stress granule disassembly in a context-specific manner. Science. 2021 June 25;372(6549):eabf6548.

15. Tolay N, Buchberger A. Comparative profiling of stress granule clearance reveals differential contributions of the ubiquitin system. Life Sci Alliance [Internet]. 2021 May;4(5). Available from: 10.26508/lsa.202000927

16. Yang C, Wang Z, Kang Y, Yi Q, Wang T, Bai Y, et al. Stress granule homeostasis is modulated by TRIM21-mediated ubiquitination of G3BP1 and autophagy-dependent elimination of stress granules. Autophagy. 2023 July;19(7):1934–51.

17. Pan CR, Knutson SD, Huth SW, MacMillan DWC. µMap proximity labeling in living cells reveals stress granule disassembly mechanisms. Nat Chem Biol. 2025 Apr;21(4):490–500.

18. Tolay N, Buchberger A. Role of the ubiquitin system in stress granule metabolism. Int J Mol Sci. 2022 Mar 26;23(7):3624.

19. Kedersha N, Panas MD, Achorn CA, Lyons S, Tisdale S, Hickman T, et al. G3BP-Caprin1- USP10 complexes mediate stress granule condensation and associate with 40S subunits. J Cell Biol. 2016 Mar 28;212(7):845–60.

20. Clague MJ, Heride C, Urbé S. The demographics of the ubiquitin system. Trends Cell Biol. 2015 July;25(7):417–26.

21. Chernov KG, Barbet A, Hamon L, Ovchinnikov LP, Curmi PA, Pastré D. Role of microtubules in stress granule assembly. J Biol Chem. 2009 Dec;284(52):36569–80.

22. Hu S, Zhang Y, Yi Q, Yang C, Liu Y, Bai Y. Time-resolved proteomic profiling reveals compositional and functional transitions across the stress granule life cycle. Nat Commun. 2023 Nov 27;14(1):7782.

23. Van Treeck B, Parker R. Principles of stress granules revealed by imaging approaches. Cold Spring Harb Perspect Biol. 2019 Feb 1;11(2):a033068.

24. Wheeler JR, Matheny T, Jain S, Abrisch R, Parker R. Distinct stages in stress granule assembly and disassembly. Elife [Internet]. 2016 Sept 7;5. Available from: 10.7554/elife.18413

25. Qian S-B, Princiotta MF, Bennink JR, Yewdell JW. Characterization of rapidly degraded polypeptides in mammalian cells reveals a novel layer of nascent protein quality control. J Biol Chem. 2006 Jan 6;281(1):392–400.

